# Molecular mimicry of plant cell-surface immune receptors by fungal secreted leucine-rich repeat proteins

**DOI:** 10.1101/2025.08.22.671746

**Authors:** Haider Ali, Elisha Thynne, Toby Ellis, Catherine Booth, Ruth Bates, Matthew S. Hope, Emma J. Wallington, Emma E. Crean, Leonhard Pachinger, Isabel M.L. Saur, Eva H. Stukenbrock, Graeme J. Kettles

## Abstract

- Leucine-rich repeat (LRR) receptor-like kinases (LRR-RLKs) are important plant immunity proteins. The wheat pathogen *Zymoseptoria tritici* produces many virulence effectors during infection; however, most remain uncharacterised. We identified a secreted protein from *Z. tritici* (ZtLRR) that consists of a single LRR domain and hypothesised that it mimics host LRR-RLKs to suppress plant immunity.
- We used transient expression to probe ZtLRR function related to production of reactive oxygen species (ROS) and cell death, two important immune processes. We used AlphaFold structural predictions with targeted yeast two-hybrid to demonstrate protein-protein interactions. Transgenic wheat allowed assessment of effector function in the natural host.
- ZtLRR suppressed ROS production and cell death in *N. benthamiana* with high potency. Structural predictions suggested high similarity to several plant LRR-RLKs and interaction with TaBAK1 was confirmed by targeted yeast two-hybrid. Transgenic wheat expressing ZtLRR was impaired in PAMP-triggered immunity (PTI) and had increased susceptibility to infection with compatible *Z. tritici*. We identified structural orthologs of ZtLRR in diverse fungal lineages and demonstrated that several of these proteins have similar immune-suppressing properties as ZtLRR.
- Our work demonstrates that molecular mimicry of host LRR-RLKs by phytopathogen sLRR effectors is effective at disrupting host immune pathways.

## Introduction

Plants use a diverse array of immune sensors to monitor for signs of pathogen invasion. These sensors can be broadly divided into intracellular and cell-surface receptors that (i) directly bind to pathogen-derived molecules, (ii) detect host-derived cues that are indicative of damage, or (iii) guard key signalling hubs that are often targeted by pathogens. To overcome this defence, many pathogens use small secreted proteins (effectors) to manipulate host defensive and metabolic processes for their own benefit. Effectors can function either intra- or extracellularly to suppress immune signalling, ensure access to nutrition or function in a myriad of other capacities that make the host more amenable to infection (Franceschetti *et al*., 2017; Giraldo and Valent, 2013; Cai *et al.,* 2023). Further, effectors rarely act in isolation but are part of an effector milieu introduced into host tissues concurrently.

Cell-surface receptors play important roles in regulation of growth and development, abiotic stress tolerance and in the earliest stages of immune responses (Tang *et al.,* 2017; Ye *et al.,* 2017; He *et al.,* 2018; Kanyuka and Rudd, 2019). Among these are the leucine-rich repeat (LRR) receptor-like kinases (LRR-RLKs) which are modular proteins featuring an extracellular LRR domain, a transmembrane domain an intracellular kinase domain. The extracellular LRR domain allows recognition of specific ligands, which may be pathogen-derived molecules such as pathogen-associated molecular patterns (PAMPs), apoplastic pathogen effectors, host-derived damage-associated molecular patterns (DAMPs) or signalling peptides (Tang *et al.,* 2017). Ligand perception by extracellular LRR domains also facilitate protein-protein interactions with other host co-receptors to form active immune complexes capable of initiating downstream immune signalling. After recognition of their specific ligand in the apoplast, LRR-RLKs frequently associate with, and signal through, a conserved partner RLK known as BRASSINOSTEROID INSENSITIVE 1-associated kinase (BAK1), which is highly conserved among monocot and dicot species (Schwessinger *et al.,* 2011; *Sun et al.,* 2013). Accordingly, heterologous expression of BAK1-dependent RLKs is possible between more distantly related species and functionality can remain intact (Zipfel *et al.,* 2006; Lacombe *et al.,* 2010; Hao *et al.,* 2016; Mendes *et al.,* 2010). Following ligand recognition and immune complex formation between the LRR-RLK and BAK1, there is initiation of an intracellular signalling cascade and pathogen countermeasures which are collectively termed PAMP-triggered immunity (PTI) or effector-triggered immunity (ETI) depending on the inducing ligand. Due to the importance and conservation of LRR-RLK/BAK1 signalling, this immune complex and its downstream signalling partners are common targets of pathogen effector attack (Shan *et al.,* 2008; Lu *et al.,* 2010; Irieda *et al*., 2019).

One evolutionary strategy used by different phytopathogens and pests to gain advantage is to employ molecular mimics. These are molecules whose function is to induce an inappropriate response in the plant through replicating the molecular structure of an endogenous host target. These can be hormone mimics, such as the *Pseudomonas* toxin coronatine, which mimics the structure of the plant hormone jasmonic acid isoleucine to trigger inappropriate stomal opening and permit pathogen entry into leaves (Zheng *et al.,* 2012; Mittal and Davis, 1995). Pathogenic *Meloidogyne spp.* And *Rotylechulus spp.* nematodes produce plant-like, C-terminally encoded peptides (CEPs) and CLAVATA3 (CLV3)/EMBRYO SURROUNDING REGION (CLE peptides) during root colonisation (Bobay *et al.,* 2013; Eves-Van Den Akker *et al.,* 2016; Gheysen and Mitchum, 2019). Endogenous CEPs and CLE peptides assist in regulating root and shoot development in response to hormonal signalling cues (Fletcher, 2020; Yamaguchi *et al.,* 2016; Kondo *et al.,* 2011; Aggarwal *et al.,* 2020; Delay *et al.,* 2013). Both classes interact with specific LRR domain-containing RLK partners (Narasimhan and Simon, 2022; Trotochaud *et al.,* 1999; Tabata *et al.,* 2014; Müller *et al.,* 2008; Rzemieniewski *et al.,* 2025), which are exploited as susceptibility factors by root knot nematodes (RKNs) producing homologous peptides (Nakagami *et al.,* 2023). Fungi can also exploit plant receptors by using mimics. Rapid alkalinisation factor (RALF) peptides are produced by plants to regulate RLK signalling via interaction with a secondary protein component of the RLK-BAK1 complex called FERONIA (Haruta *et al.,* 2014; Stegmann *et al.,* 2017). FERONIA is a malectin-like kinase that assists in scaffolding RLKs to BAK1, thereby enabling association and signalling (Stegmann *et al*., 2017). Several fungi (and RKNs (Zhang *et al.,* 2020)) produce RALF peptides that can hijack FERONIA activity (Masachis *et al*., 2016; Thynne *et al*., 2017; Liao *et al*., 2023) to enhance host susceptibility. Combined, the examples of CEP, CLE and RALF peptides are all example of plant-like protein mimics produced by pathogens to target LRR-RLK signalling.

*Zymoseptoria tritici* is a hemibiotrophic ascomycete fungus which causes the wheat disease Septoria tritici blotch (STB) (Kettles and Kanyuka, 2016). This fungus grows epiphytically on wheat leaves before invading through open stomata and growing through the apoplastic space of the mesophyll (Fantozzi *et al.,* 2021). This apoplastic colonisation is asymptomatic, with the fungus able to avoid triggering host immune responses for an extended period. Following asymptomatic growth, *Z. tritici* infections transition to necrotrophy, with appearance of disease symptoms and death of host cells. *Z. tritici* secretes effectors during infection, which can display stage-specific expression patterns (Mirzadi Gohari *et al.,* 2015; Rudd *et al*., 2015; Haueisen *et al.,* 2019; Battache *et al*., 2022; Haueisen *et al.,* 2025; Alassimone *et al.,* 2024). Some effectors are highly expressed during the asymptomatic growth phase such as the LysM domain-containing effectors (Marshall *et al.,* 2011; Sánchez-Vallet *et al*. 2020; Tian *et al*. 2021), which inhibit chitin-triggered defences. Others, such as the toxic ribonuclease Zt6, peak in expression later in infection at the transition to necrotrophy (Kettles *et al.,* 2018). We recently described a large group of *Z. tritici* effectors with immune-suppressing phenotypes in plants (Thynne *et al.,* 2024). Many of these effectors were small proteins with no obvious domains or motifs in their primary sequence and poor structural predictions which makes further characterisation challenging. In parallel, we identified an unusual effector that was highly expressed during symptomless colonisation that contained a single LRR domain. Given the important role of LRR domain-containing proteins in plant immune receptors, we speculated that this effector might have an important role in immune suppression during early colonisation.

Here we report that the secreted LRR (sLRR) effector from *Z. tritici* (ZtLRR) functions as a potent inhibitor of immune responses in both *N. benthamiana* and wheat (*Triticum aestivum*). We provide evidence that it does this through a novel form of molecular mimicry that targets the apoplastic regions of cell-surface immune complexes involving host LRR-RLKs.

## Materials and methods

### Plant materials and growth conditions

*N. benthamiana* plants were used for all *Agrobacterium*-mediated transient expression assays. Seeds were sown in peat-free compost (Jiffy Products International, Moerdijk, Netherlands), consisting of a 3:1 mixture of peat-free potting substrate and Silvaperl perlite, and maintained in controlled-environment growth chambers under long-day conditions (16 h light/8 h dark) at a constant temperature of 22°C and ∼60% relative humidity. Plants were watered every three days. All experiments were conducted using fully expanded leaves of 5–6 week-old plants.

For infection assays and reactive oxygen species (ROS) burst measurements in wheat, *Triticum aestivum* cultivar ‘Cadenza’ and transgenic lines expressing ZtLRR effector were used. Wheat seeds were germinated on moist filter paper in the dark at room temperature for 2-3 days before being transferred to 22.5 × 17.5 cm half-trays. Plants were grown in a controlled-environment glasshouse at 20–22°C with a 16 h light/8 h dark photoperiod and ambient humidity.

### Generation of wheat transgenics

For expression of full-length *ZtLRR* (including native signal peptide) in wheat, PCR was used to (i) introduce a ribosome binding site (CCACC) immediately upstream of the *ZtLRR* start codon, and (ii) reintroduce a stop codon, using pEAQ-HT-DEST3(*ZtLRR*) plasmid (Kettles *et al*., 2017) as template. This modified sequence was recombined by Gateway BP reaction into the pDONR207 vector. The coding sequence was then transferred by Gateway LR reaction into the binary vector pEW472-Ubi-R1R2 to create pMSH83 for expression under control of the *Zea mays* ZmUbi promoter for high-level constitutive expression in wheat. Immature embryos isolated from wheat variety Cadenza were transformed by inoculation with *Agrobacterium tumefaciens* strain LBA4404 pSB1 containing pMSH83 essentially as described previously but with a 2 or 3-day co-cultivation period and 2mg/l 2,4-D in the CO1 medium (Milner *et al*., 2024). Thirty-nine regenerated plants with good root development were transferred to Jiffy 7 pellets and acclimatised in a propagator. DNA from the transgenic lines was extracted from leaf material and analysed for nptII copy number (Milner *et al.,* 2024). Following identification of lines with single-copy transgene insertions in T0 seedlings, RNA was extracted from leaves of 4–5-week-old T0 plants using Trizol and cDNA synthesised using Superscript III reverse transcriptase (Thermo Fisher Scientific) following the manufacturer’s instructions. qRT-PCR was performed using a previously described method (Kettles et al. 2019). Expression of *ZtLRR* was quantified by the 2^-ΔCt^ method relative to a panel of three internal reference genes (*Ta-CDC48*, *Ta-β-tubulin* and *Ta-Ubiquitin*) to identify T0 lines with the highest *ZtLRR* expression level. Individual plants of these lines were self-pollinated and T1 seed collected. T1 plants were grown in the same conditions and leaves of seedlings harvested for genomic DNA extraction. qPCR was used to discriminate transgene copy number in homozygotes, heterozygotes and null segregants. Homozygous and null segregants were retained and self-pollinated to produce T2 seed for plant assays and further bulking to T3 seed.

### Fungal isolates, culture conditions, and wheat infection assays

*Z. tritici* isolates IPO323 and IPO88004 were used in this study as described previously (Ali *et al.,* 2024). Briefly, cultures were maintained on yeast peptone dextrose agar (YPDA; 2% glucose, 2% peptone, 1% yeast extract, and 1.5% agar; Formedium, Kings Lynn, UK) and incubated at 18°C in the dark for 5–6 days. To prepare spore suspensions for infection, fungal material was harvested directly from the surface of YPDA plates using a sterile inoculating loop and suspended in sterile distilled water containing 0.01% (v/v) Tween-20. The suspension was vortexed thoroughly to disperse the spores and adjusted to a final concentration of 1 × 10⁶ spores mL⁻¹ using a haemocytometer.

Wheat plants at the second-leaf stage were inoculated by applying the spore suspension to the adaxial surface of the second leaf using sterile cotton swabs. Immediately after inoculation, the plants were enclosed in sealed transparent propagation boxes (Garland Products Ltd., UK) to maintain high humidity and incubated under these conditions for 48 h. Plants were then returned to standard glasshouse conditions and maintained under a 16 h light/8 h dark photoperiod at 20– 22°C until disease assessment at 21 days post-inoculation (dpi).

Disease progression was quantified by measuring the green leaf area (GLA) from high-resolution images of infected leaves processed using MIPAR image analysis software (MIPAR Software LLC, USA)(Qutb BMC Plant Biology 2024). The extent of necrotic and chlorotic lesions was used as a proxy for symptom severity. To assess asexual pycnidiospore production, infected leaves were harvested at 21 dpi, submerged in sterile distilled water, and vortexed vigorously for 2 min. Pycnidiospore concentrations were determined using a haemocytometer under a light microscope. Each treatment included a minimum of three biological replicates per treatment.

### Yeast two-hybrid

Yeast two hybrid (Y2H) assays were performed using the MatchMaker Gold Y2H System (Takara Bio). From this, we used the empty pGBKT7 (binding domain) and empty pGADT7 (activation domain) vectors. Additionally, we used the kit control plasmids: pGADT7-T-protein; pGBKT7-p53 (T-protein interactor); and pGBKT7-Lam (T-protein non-interactor). TaBAK1 (NCBI accession: CFC21_109359) extracellular domain (LRR) (aa N23-G210) and intracellular domain (kinase) (aa R235-R597) encoding DNA were each independently synthesised into the pGADT7 plasmid (TWIST Biosciences). ZtLRR encoding-DNA (without signal peptide) was synthesised into the pGBKT7 plasmid (TWIST Biosciences). Combinations of pGADT7 and pGBKT7 plasmids were co-transformed into yeast strain Y2H Gold, using the Yeastmaker Yeast Transformation System 2 (Takara Bio). Transformants were selected on synthetic defined medium (SD-M), lacking both leucine (-L) and tryptophan (-W). Yeast colonies were picked with a pipette tip and diluted in sterile water. 5µl of yeast samples were plated on both SD-M (-L,-W) and SD-M(-L,-W,-H,-A + x-alpha-gal). Colonies were left to grow for three days at 28°C before visualisation.

### Identification of sLRR orthologs in other fungi

To identify functional orthologs of ZtLRR, we performed BLASTp searches using the full-length ZtLRR amino acid sequence against the NCBI non-redundant protein sequences (nr) database. We selected 15 candidate sLRR proteins from fungal species representing different taxonomic groups, host range and lifestyle (necrotrophic pathogen, hemibiotrophic pathogen, saprotroph, endophyte). Candidate sLRRs were confirmed as secreted proteins and/or likely effectors using the tools SignalP/TargetP for conventional secretion, EffectorP for possible effector activity and SecretomeP/OutCyte for unconventional secretion. Candidates predicted to be secreted via classical or unconventional pathways were retained, regardless of the percentage identity or e-value. Codon-optimised sLRR sequences were synthesised directly into the pTwist (Chlor) Gateway entry vector (Twist Biosciences, San Francisco, USA). Following plasmid isolation (QIAprep Spin Miniprep, Qiagen), plasmids were linearised with *NsiI* (NEB) and recombined into the pEAQ-HT-DEST3 plant expression vector using LR Clonase II (Thermo Fisher Scientific). The final constructs were sequence-verified (Eurofins Genomics) and transformed into *A. tumefaciens* GV3101 (pMP90) by electroporation.

### Agroinfiltration and transient expression in *N. benthamiana*

*Agrobacterium tumefaciens* GV3101 (pMP90) carrying *ZtLRR*, *Zt6*, *Zt12* or *sGFP* in the pEAQ-HT-DEST3 vector were generated previously (Kettles *et al.,* 2017; Thynne *et al.,* 2024). The *ZtLRRΔSP* construct was generated in this study by removing the sequence encoding the first 19 amino acid secretion signal peptide of *ZtLRR* by PCR and subcloning into the same vector. Agrobacterium strains were cultured shaking (220rpm) for two nights at 28°C in Luria–Bertani (LB) medium supplemented with kanamycin (50 µg/mL⁻¹) and gentamicin (25 µg/mL⁻¹). Cells were harvested by centrifugation at 13,000 rpm for 1 min, repeated twice, and then washed twice with infiltration buffer (10 mM MgCl₂, 10 mM MES, pH 5.6). The final pellet was resuspended in the same buffer to an optical density of OD_600_=1.2 and supplemented with 150 µM acetosyringone. The suspensions were incubated at room temperature for 2–3 h in the dark to induce virulence gene expression. Suspensions were pressure-infiltrated into the abaxial surface of fully expanded leaves from 5–6-week-old *N. benthamiana* plants using a 1 mL needleless syringe. Leaf tissue was harvested 2–3 dpi for ROS burst assays and 5 dpi for cell death suppression assays. To quantify cell death, autofluorescence was measured using an Amersham Typhoon scanner (Cytiva) following the method of Xi and coworkers (Xi *et al.,* 2021). 15-20 leaves were used (two leaves/plant) in each experiment.

### ROS burst assays in *N. benthamiana* and wheat

ROS burst assays were performed using *N. benthamiana* and *T. aestivum* (cv. Cadenza and ZtLRR transgenic lines) using a previously described chemiluminescence-based method. For *N. benthamiana*, leaves from 5–6-week-old plants were agroinfiltrated, as described above. At 2–3 dpi, eight leaf discs (3 × 3 mm) were excised from each infiltration site using a sterile scalpel and floated overnight in 200 µL sterile distilled water in white 96-well microplates (Thermo-Fisher Scientific, UK) to reduce the background signal and wounding effects. The following day, the water was replaced with 150 µL of elicitation buffer containing horseradish peroxidase (HRP; 20 ng mL⁻¹), the luminol derivative L-012 (20 µM; Wako Chemicals), and one of the following elicitors: 100 nM flg22, 100 µg mL⁻¹ chitin (Megazyme, Ireland), or 100 µg mL⁻¹ laminarin (Sigma-Aldrich). Luminescence was measured immediately using a PHERAstar FS plate reader (BMG Labtech, Germany) for approximately 2 h (90 cycles). Each treatment included at least eight biological replicates and each assay was performed thrice.

For wheat, leaf strips (∼5 cm) were harvested from the second leaf of 2–3-week-old wheat seedlings. The strips were cut into ∼5 mm long strips using a sterile scalpel and floated overnight in sterile distilled water in a white 96-well plate. The next day, ROS production was triggered by replacing the water with 150 µL of elicitation buffer containing 1 mg mL⁻¹ chitin, 1 mg mL⁻¹ laminarin, or 50 µM flg22, supplemented with HRP and L-012 at the same concentrations as above. Luminescence was measured in real time for 2 h (90 cycles) using a PHERAstar FS microplate reader. Each genotype (Cadenza or ZtLRR transgenic line) was tested using three independent plants, with a minimum of two leaf strips per well. The experiment was performed twice. ROS output was quantified as relative light units (RLU) to assess the total oxidative burst for each treatment. Statistical analysis was conducted in RStudio using one-way ANOVA, followed by Tukey’s HSD test.

### ROS burst assays in wheat protoplasts

Protoplast isolation and transfection were performed as described (Saur et al. 2019) with the following deviations. Briefly, protoplasts were extracted from the first leaf of 7-10 day old seedlings of cv. Fielder. For each construct (*pZmUbi:GFP* and *pZmUbi:ZtLRR*), 1.5 ml protoplasts at a concentration of 3.5 x 10^5^ cells/ml were transfected with 150 µg of plasmid. Following washing, 200 µl of protoplasts from each transfection were resuspended in 0.5 ml regeneration buffer and divided as 50 µl aliquot/well to white 96-well plates. The plates were stored overnight in the dark at 18 °C and ROS was measured 14-20 hours later. For this, to each well containing transfected protoplasts, 50 µl of regeneration buffer supplemented with 20 µg/ml HRP, and 20 µM L-012 luminol, and either without elicitor (mock) or with elicitor (400 nM flg22,20 µg/ml chitin (NA-COS-Y) was added (Mahdi et al. 2020). This resulted in final concentrations of 10 µg/ml HRP, 10 µM L-012, 200 nM flg22 and 10 µg/ml chitin. Luminescence was measured for 1 h with 1 sec measurement/well using a TECAN plate reader. Four samples per elicitor (flg22, chitin, mock) were measured for the respective transfected protoplast constructs. The experiment was performed three times in total using protoplasts derived from independently grown plants each time.

## Results

### ZtLRR suppresses cell death and ROS burst immune responses when directed to the apoplast in *N. benthamiana*

In a recent study, we demonstrated that numerous *Z. tritici* candidate effectors can suppress plant immune responses such as effector-induced cell death and the PAMP-induced ROS burst when expressed in *N. benthamiana* (Thynne *et al*., 2024). To test whether a *Z. tritici* sLRR-domain containing protein (ZtLRR, previously annotated as ZtIPO323_123670, Mycgr3_97371, evm.TU.chr_13.171) could induced similar phenotypes, we used *Agrobacterium*-mediated transient expression (Agroexpression) to express *ZtLRR* in leaves of *N. benthamiana. ZtLRR* was previously cloned into an Agroexpression vector with its native N-terminal signal peptide for secretion to the leaf apoplastic space (Kettles *et al.,* 2017). As a negative control, we expressed *GFP* fused to the *Nicotiana tabacum PR1a* secretion signal peptide (*sGFP*) (Thynne *et al.,* 2024). First, we assessed the ability of ZtLRR to suppress cell death triggered by the *Z. tritici* effectors Zt12, which induces large-scale host transcriptional reprogramming and BAK1/SOBIR1-dependent cell death in *N. benthamiana* (Kettles *et al.,* 2017; Welch *et al*., 2022) and Zt6, a ribonuclease toxin that induces BAK1/SOBIR1-independent cell death (Kettles *et al*., 2018). Both *Zt6* and *Zt12* induced obvious macroscopic cell death when co-expressed with *sGFP* at three days post-infiltration (3 dpi)(Fig.1a). Similarly, *Zt6* induced cell death when co-expressed with *ZtLRR*. However, co-expression of *Zt12* with *ZtLRR*, resulted in less obvious cell death (Fig.1a). To confirm cell death suppression, agroinfiltrated leaves were scanned by fluorescence imaging (Xi *et al.,* 2021) to quantify the relative reduction in leaf autofluorescence caused by the dead/dying cells present in each leaf patch. In leaf tissue with widespread cell death, autofluorescence is decreased, whilst in living tissue it remains high. There was no significant difference in autofluorescence intensity between the *Zt6*-expressing samples, indicating no quantitative suppression of Zt6-triggered cell death. However, autofluorescence was significantly increased in the ZtLRR+Zt12 treatment compared to the Zt12+sGFP control, thus confirming visual observation of suppression of Zt12-triggered cell death (Fig.1a).

**Figure 1.**
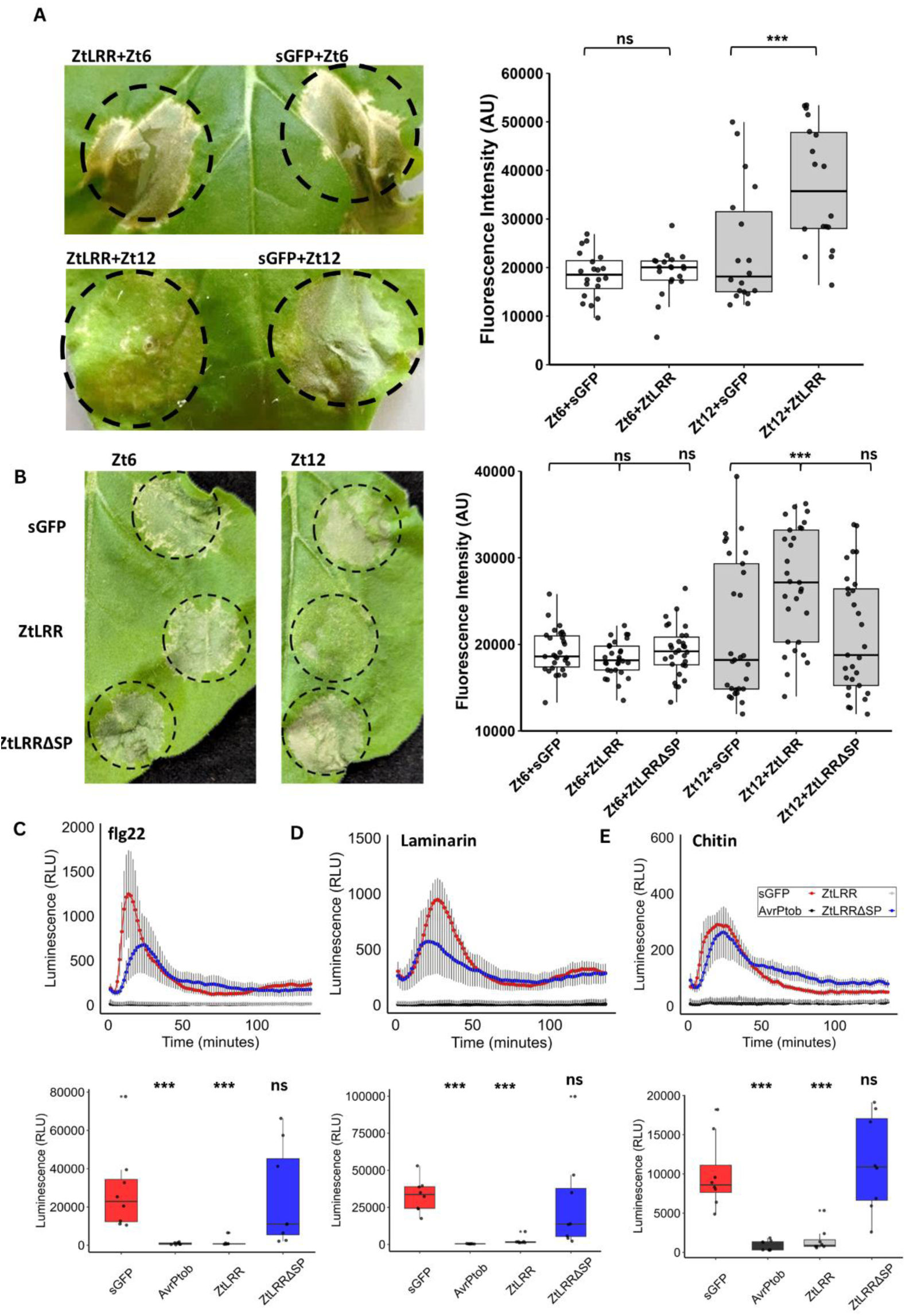
ZtLRR suppresses effector-triggered cell death and PAMP-triggered ROS production in *N. benthamiana*. (A) ZtLRR suppresses cell death triggered by Zt12, but not Zt6, when co-expressed in *N. benthamiana*. Leaves photographed at 5 days post-infiltration (dpi). Cell death was quantified by red fluorescence imaging (arbitrary units, AU) from 20 leaves per treatment. White and grey boxes represent Zt6 and Zt12 treatments respectively. Asterisks indicate statistically significant differences at \**p*<0.05, \*\**p*<0.01, and \*\*\**p*<0.001; ns, not significant, as performed by Student’s t-test. (B) Deletion of the ZtLRR signal peptide (ZtLRRΔSP) abolishes suppression of Zt12-induced cell death. Experiments performed as described above. Statistical differences were assessed using Tukey’s HSD test (*P* < 0.001; ns, not significant). (C-E) Apoplastic-dependent suppression of PAMP-induced ROS production induced by flg22 (C), laminarin (D), or chitin (E). sGFP (red) and AvrPtoB (dark grey) were used as negative and positive controls. Real-time ROS production shown in upper panels, cumulative ROS in lower panels. ROS levels presented as relative luminescence units (RLU) over time. Asterisks indicate statistically significant differences from the sGFP control at \**p*<0.001; ns, not significant, as performed by Tukey’s HSD test.

We hypothesised that similar to other characterised *Z. tritici* effectors, the immune-suppressing activity of ZtLRR may be dependent on localisation to the apoplastic environment. To test this, we recloned *ZtLRR* without its signal peptide (*ZtLRRΔSP*) and agroexpressed this construct alongside full-length *ZtLRR* in Zt6/Zt12-triggered cell death assays. Removing the signal peptide from *ZtLRR* had no impact on Zt6-triggered cell death as expected (Fig.2b). However, in leaf patches co-expressing *Zt12* and *ZtLRRΔSP* there was consistent and strong cell death similar to the Zt12+sGFP control, indicating a loss of cell death suppression. In agreement, autofluorescence intensity in Zt12+ZtLRRΔSP samples returned to that of the sGFP control and was significantly lower than for ZtLRR directed to the apoplast (Fig.1b). These observations indicate that ZtLRR is required to be localised to the apoplast for its cell death-suppressing activity.

**Figure 2.**
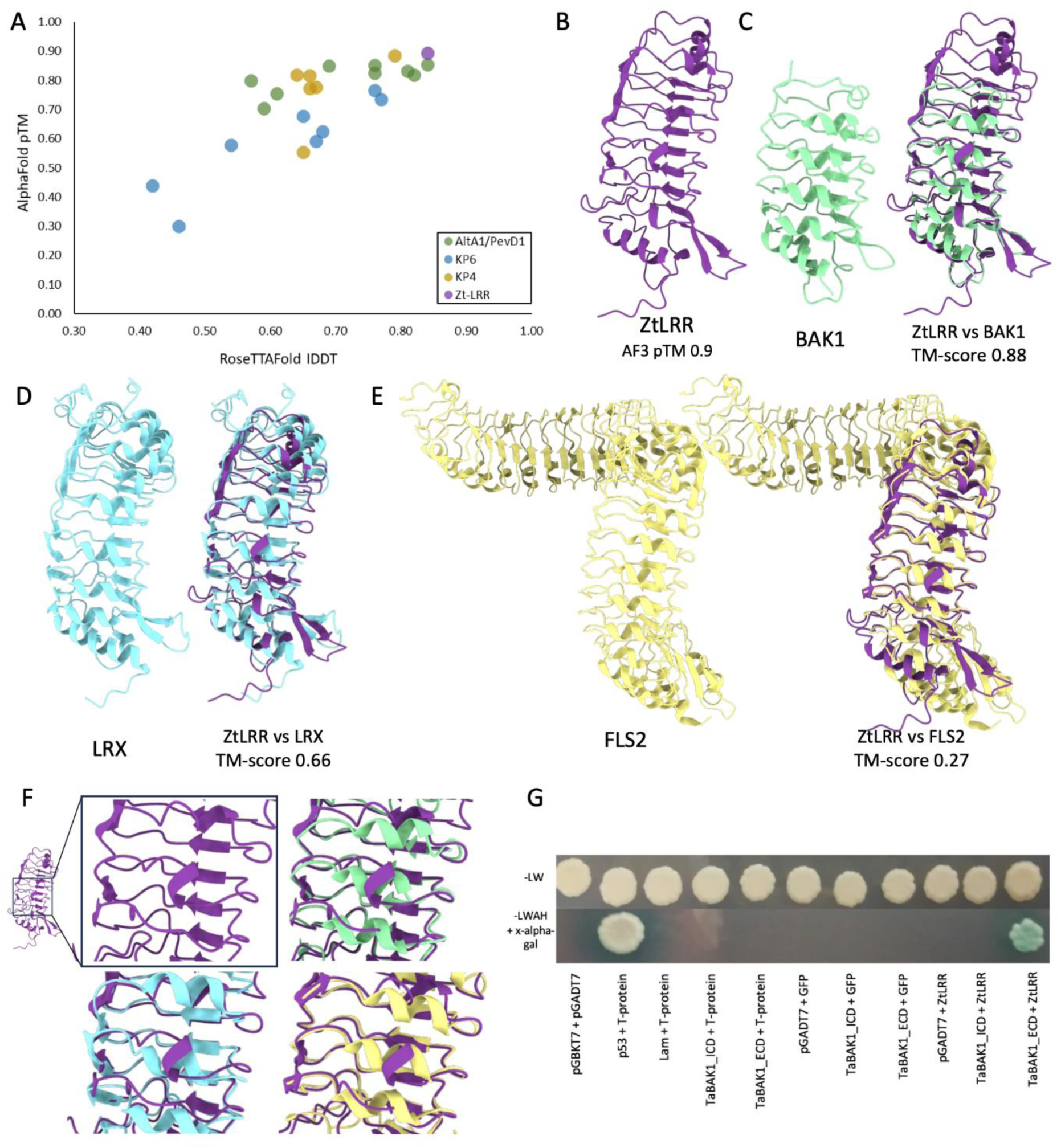
ZtLRR is a predicted structural homologue of plant LRR-RLKs and can interact with TaBAK1. A) Plot describing the structural prediction confidence statistics of *Z. tritici* effectors predicted with AlphaFold2 (y-axis; pTM) and RoseTTAFOLD (x-axis; IDDT). The structural prediction statistics for three other families of effectors, KP6, KP4, and Alt-a1, were included for comparison to ZtLRR, whose structure was predicted with the highest confidence. B) The structure of ZtLRR (purple) predicted with AlphaFold3. C) The solved structure of BAK1 (from rice; green) both alone and aligned to ZtLRR (purple). D) The solved structure of LRX (from Arabidopsis; cyan) both alone and aligned to ZtLRR (purple). E) The solved structure of FLS2 (from Arabidopsis; yellow) both alone and aligned to ZtLRR (purple). F) Enlarged images of the central three LRR units of ZtLRR (purple) and side-by-side comparisons with its alignments to the corresponding LRR units from BAK1 (green), LRX (cyan), and FLS2 (yellow) to demonstrate similarity. G) Yeast two-hybrid interaction between ZtLRR and wheat BAK1 (NCBI accession: CFC21_109359) intracellular (TaBAK1-ICD aa: R235-R597) and extracellular domains (TaBAK1-ECD aa: N23-G210), plated on SD-M(-LW) and SD-M(-LWAH + x-alpha-gal). Only the positive control plasmids (pGBBKT7-p53 and pGADT7-T-protein) and ZtLRR with TaBAK1-ECD enabled yeast growth on the interaction selection SD-M(-LWAH + x-alpha-gal), indicating a putative interaction between ZtLRR and the extracellular LRR domain of BAK1.

In our previous study, we demonstrated that suppression of the PAMP-induced ROS burst is a common feature of *Z. tritici* effectors (Thynne *et al*., 2024). To test if this was also the case for ZtLRR, we expressed either full-length or intracellular ZtLRR alongside a sGFP control and triggered ROS production by supplying the bacterial and fungal PAMPs flg22, laminarin, and chitin. We also expressed the *Pseudomonas syringae* effector *AvrPtoB* as a positive control for ROS burst suppression. Leaf discs were taken from plants expressing *sGFP*, *ZtLRR*, *ZtLRRΔSP* or *AvrPtoB* and treated with flg22, laminarin or chitin in a luminol-based 96-well plate assay. In comparison to the *sGFP* control, ROS production was almost completely abolished in tissue expressing either *ZtLRR* or *AvrPtoB*, for each of the three PAMP treatments (Fig.1c-e). In contrast, ROS bursts in tissue expressing *ZtLRRΔSP* were returned to similar levels as the *sGFP* controls (Fig.1c-e). Although the luminescence maxima for *ZtLRRΔSP* was lower than *sGFP* during flg22 and laminarin treatment, there were no statistically significant differences in cumulative ROS production. These findings demonstrate that ZtLRR can suppress PAMP-induced ROS bursts for flg22, laminarin, and chitin at a similar potency to AvrPtoB, and that this suppressive activity requires ZtLRR localisation to the apoplast.

### ZtLRR has predicted structural homology to plant cell-surface RLKs and physically interacts with the co-receptor TaBAK1

We previously demonstrated immune-suppressing activity of several *Z. tritici* candidate effectors. However, many of these proteins have no obvious domains or motifs at the primary sequence level, lack experimentally validated structures or have poor structural predictions from tools such as AlphaFold. Therefore, understanding the mechanism(s) of effector function is challenging. However, as ZtLRR contains an obvious LRR domain (InterProScan (Jones *et al.,* 2014)) from its primary sequence, we speculated that Alphafold would generate a high-confidence structural model. We made structural predictions for ZtLRR and a number of other *Z. tritici* effectors using RoseTTAFold (Baek et al. 2021) and AlphaFold2 (Jumper et al. 2021). We compared the AlphaFold2 pTM and RoseTTAFold IDDT prediction probability scores for ZtLRR to those of other *Z. tritici* effectors, from three structural families that have been partly characterised; the killer-like protein four (KP4), killer-like protein six (KP6); and PevD1/AltA1 families (Fig.2a). We included these as references as they belong to structural families that are enriched amongst plant pathogens including *Z. tritici* (Seong and Krasileva, 2023). Further, there is a solved crystal structure for a KP6 effector from *Z. tritici* (De Guillen *et al.,* 2024) and for an AltA1 effector from *Alternaria alternata* (Chruszcz *et al*., 2012). Compared to effectors in these families, ZtLRR had the highest structural prediction probability score (AlphaFold2 pTM of 0.895 and RoseTTAFOLD IDDT of 0.84) (Fig.2a,b) suggesting it may be possible to infer function from such a high-confidence prediction. We then used this model (Fig.2b) for a structural homology search using the FoldSeek database (van Kempen *et al*., 2024). This generated many high-confidence structural matches, with the most similar being the ectodomains of plant LRR-RLKs rather than other microbial LRRs. From among these hits were the conserved LRR-RLK co-receptor BAK1 (SERK3), the flg22-interacting LRR-RLK FLS2 and the plant LRR protein LRX, which assists in regulation of RLK/BAK1/FERONIA complex formation. The extracellular LRR domains (ectodomains) for these three proteins were extracted from their experimentally validated structures (PDB codes 4Q3G (McAndrew *et al*., 2014), 4MN8 (Sun *et al*., 2013), 6QXP (Moussu *et al*., 2020)) and overlaid against the ZtLRR model (Fig.2c-f). Given the high structural similarity between ZtLRR and several LRR-RLKs, we hypothesised that this may be indicative of the mechanism of immune suppression observed in our cell death and ROS burst assays (Fig.1). Zt12-triggered cell death and the flg22-triggered ROS burst both require the BAK1 co-receptor for immune signalling. We speculated that mimicry of an LRR-RLK ectodomain, such as FLS2, may allow ZtLRR to directly bind and disrupt apoplastic immune complex formation. To test this, we used targeted yeast two-hybrid to co-express *ZtLRR* with the *T. aestivum BAK1* (*TaBAK1*) intracellular and extracellular domains (Fig.2g). We found no evidence of autoactivation of any of the constructs when expressed singularly. Yeast growth on selective media was only observed when *ZtLRR* and the *TaBAK1* ectodomain were expressed together. Expression of *ZtLRR* and the intracellular kinase domain of *TaBAK1* did not result in yeast growth (Fig.2g). This data is indicative of a direct interaction between ZtLRR and the TaBAK1 ectodomain in yeast cells.

### ZtLRR suppresses PTI in wheat protoplasts

Given the immune suppression exerted by ZtLRR in non-host *N. benthamiana*, we next assessed whether similar phenotypes could be observed in wheat, the natural host for *Z. tritici*. First, we expressed *ZtLRR* and *GFP* individually in protoplasts obtained from the wheat cultivar Fielder. These *ZtLRR*- and *GFP*-expressing protoplasts were then treated with flg22, a chitin analogue or mock treatment (water) and incubated with luminol and HRP to measure ROS production via chemiluminescence. For both PAMP treatments, total RLUs were significantly decreased for protoplasts expressing *ZtLRR* compared to protoplasts expressing *GFP* (Fig.3a). This indicated that, similar to our observation *N. benthamiana*, ZtLRR can suppress PAMP-triggered ROS in wheat.

**Figure 3.**
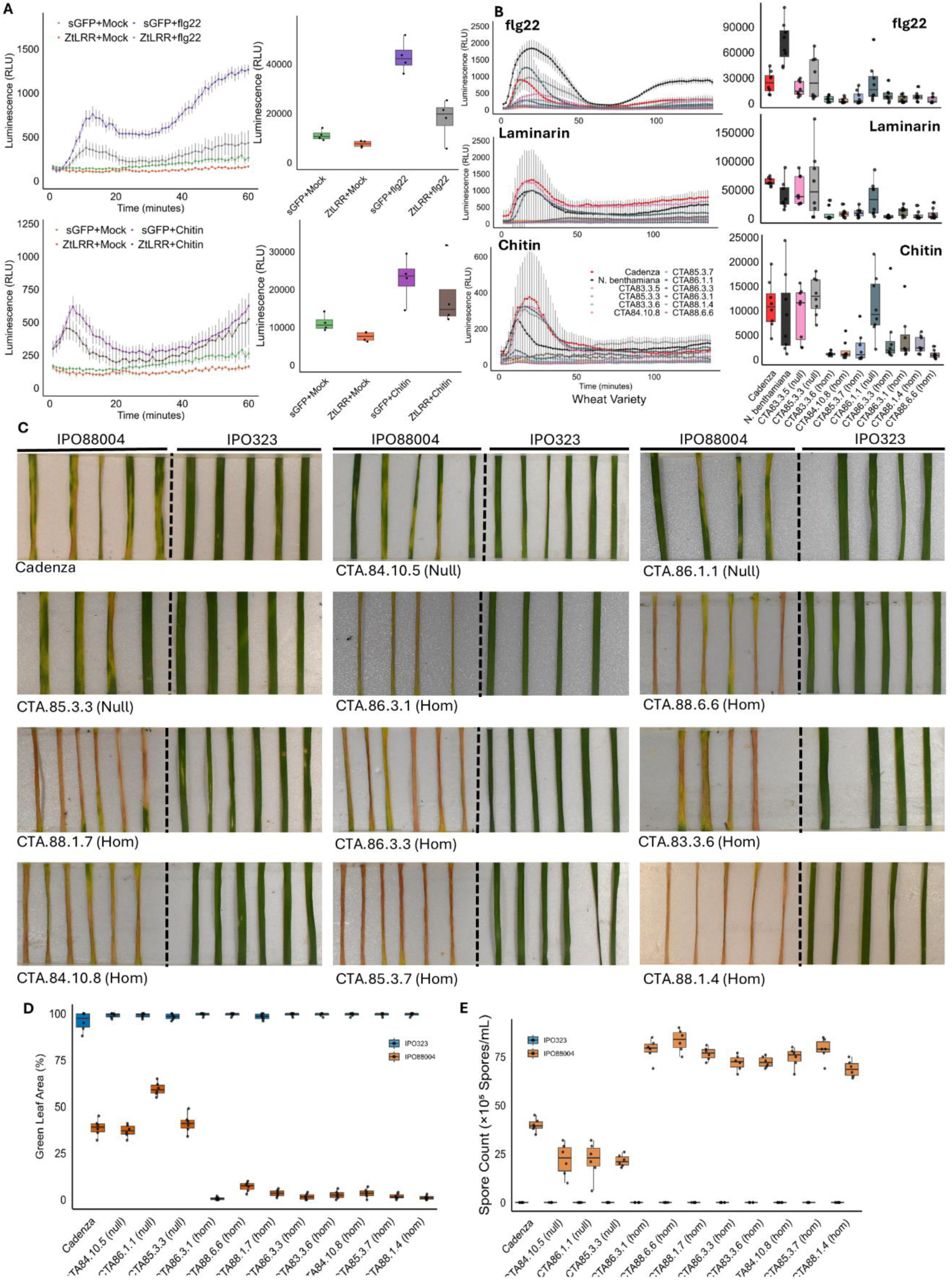
ZtLRR expression in wheat suppresses PAMP-triggered ROS production and enhances susceptibility to *Z. tritici.* (A) Expression of ZtLRR in wheat protoplasts reduces flg22- or chitin-induced ROS production in comparison to a GFP control. Protoplasts were generated from wheat cv. Fielder and treated with flg22 (200 nM) and chitin (10 µg mL⁻¹). Real-time ROS production shown in left panels, cumulative ROS over 60 min in right panels. ROS production was measured as relative luminescence units (RLU) over one hour. (B) Transgenic wheat expressing ZtLRR (homozygous, hom) showed ROS burst suppression following flg22 (50 µM), laminarin (1 mg mL⁻¹) and chitin (1 mg mL⁻¹) treatment compared to wild-type Cadenza and null segregant controls. Real-time ROS production shown in left panels, cumulative ROS in right panels. Bars represent means ± SE. (C) ZtLRR-expressing wheat (homozygous, hom) displayed enhanced disease symptoms to IPO88004 (orange) infection, but not IPO323 (blue), compared to wild-type and null segregant controls. Leaf images were taken at 21 days post-inoculation (dpi). (D) Quantification of green leaf area (GLA) at 21 dpi from images shown in (C). Bars represent the mean ± SE. (E) Increased sporulation of IPO88004 (orange) but not IPO323 (blue) on ZtLRR transgenic homozygous lines compared to wild-type and null segregants. Spore counts obtained via leaf wash at 21 dpi.

### *ZtLRR* expression in transgenic wheat suppresses ROS and enhances virulence of a compatible fungal isolate

To further explore the impact of *ZtLRR* expression in wheat, we generated stable transgenic lines expressing *ZtLRR* in the spring wheat cultivar Cadenza. Homozygous and null segregants were identified in the T2 generation, and T3 plants were used for PAMP-induced ROS assays (Fig.3b) using similar methodology as for *N. benthamiana*. In these experiments, leaves of wild-type Cadenza control plants, three T3 null segregant lines and eight lines homozygous for the *ZtLRR* transgene were exposed to flg22, laminarin and chitin with ROS production measured for ∼2 hours. For all three PAMPs, ROS levels generated in the three null segregant lines were similar to that generated in the wild-type Cadenza control leaves (Fig.3b). However, ROS levels were significantly lower in seven of eight of the homozygous transgenic lines against all three PAMPs. This indicated that similar to expression in *N. benthamiana* plants and wheat protoplasts, ZtLRR suppresses PAMP-induced ROS in whole wheat leaves.

We next made use of the same wheat lines in a pathogen bioassay using the *Z. tritici* isolates IPO323 and IPO88004. IPO323 is avirulent on Cadenza as it expresses an avirulent allele of the *AvrStb6* effector that triggers *Stb6*-mediated resistance (*Qutb et al.,* 2024). In contrast, IPO88004 has a virulent allele of *AvrStb6* which doesn’t trigger *Stb6* resistance and this isolate is fully compatible. In these infections, virulence was scored by two metrics; (i) loss of green leaf area (GLA), and (ii) asexual spore (pycnidiospore) counts from leaf washes when the experiment was terminated. In these infections, we observed we observed a more rapid loss of GLA in the homozygous lines compared to both the null segregants and wild-type Cadenza when leaves were inoculated with the compatible isolate IPO88004 (Fig.3c,d). Furthermore, spore counts in the *ZtLRR* homozygous lines were considerably higher than the wild-type and null segregants (Fig.3e). This indicates that for the compatible Cadenza-IPO88004 interaction, *ZtLRR* overexpression conferred both a more rapid development of disease symptoms (chlorosis/necrosis) and increased level of fungal asexual sporulation. No disease symptoms were observed in any leaves inoculated with IPO323, indicating that *ZtLRR* overexpression did not confer virulence to a normally avirulent fungal isolate (Fig.3c,d).

### sLRR proteins are distributed across the fungal kingdom and have immune-suppressive capacity

To investigate whether sLRR effectors may be important virulence factors for other fungi, we searched for plant-like LRR orthologs encoded in the genomes of other fungal species. ZtLRR was used as a BLASTp query to search the NCBI-NR database (e-value of 1e-5). Similar to the earlier structural analyses (Fig.2), the majority of BLASTp hits identified were not fungal, but from plants. Among the fungal BLAST hits, we identified orthologs primarily among the Dothideomycetes and Sordariomycetes (Fig.S1, File S1). The presence/absence of sLRRs does not appear to strictly correspond to species relationship. Many species or genera of fungi that are closely related to harbouring species, lack sLRR orthologs. For example, we only observed three *formae speciales* of *Fusarium oxysporum* that have sLRR orthologs. Similarly, many Dothideomycete species possess orthologs, but this is not universal. For example, we did not identify orthologs in any of the *Botryosphaeriaceae* (a family well-represented by plant pathogenic fungi). Some genera or families of fungi are enriched with sLRRs. For example, every examined member of the *Mycosphaerellaceae* (which includes *Z. tritici*) possess a sLRR-encoding gene. In addition to presence/absence of the sLRRs, we also observed variation in copy number among species. Where *Zymoseptoria spp.* each only have a single sLRR copy, *Bipolaris spp.* have multiple copies (for example, *B. sorokiniana* possesses two orthologs and *B. zeicola* has six orthologs). We then selected 15 sLRRs from a diverse range of fungi and oomycetes for functional characterisation (Table S1). Of this set, 14 were from fungal species, with 11 representing the Ascomycota, two from the Mucoromycota and one example from the Chytridiomycota. We also included a single representative of the Oomycota (*Phytophthora infestans*). When compared to ZtLRR, amino acid sequence identity ranged from 22.7% (*Bipolaris maydis*, BmLRR,) to 39.1% (*Globomyces pollinis-pini*, GppLRR). All selected examples are predicted secreted proteins, with 10 predicted to be conventionally secreted through the signal peptide-mediated ER-Golgi pathway (by SignalP/TargetP). The remaining four were predicted to be directed through the unconventional protein secretion (UPS) pathway (by SecretomeP/Outcyte)(Table S1). Each of the sLRRs can be separated into the two observed types (based on BLAST analysis); plant-like sLRRs, and bacterial-like sLRRs.

To assess whether these 15 sequence-dissimilar, but structurally conserved, fungal and oomycete LRRs could suppress plant immune responses, we agroexpressed all candidates in *N. benthamiana* and tested their ability to suppress PAMP-induced ROS bursts (Fig.4a). We found that similar to the *AvrPtoB* and *ZtLRR* positive controls, six sLRRs suppressed flg22-induced ROS, five suppressed laminarin-induced ROS and six suppressed chitin-induced ROS. Of these candidates, five (*Plenodomus lindquistii* (*PlLRR*), *Colletotrichum orbiculare* (*CoLRR*), *Rhynchosporium commune* (*RcLRR*), *Fusarium oxysporum* (*FoLRR*), *Ramularia collo-cygni* (*RccLRR*)) suppressed ROS induced by all three PAMPs, whilst one candidate (*Mucor ambiguus, MaLRR*) suppressed ROS induced by flg22 and chitin only. The remaining nine candidates did not show any ROS suppressive activity. We continued with the six candidates that demonstrated ROS suppressive activity and assessed whether they could also suppress Zt6- and Zt12-induced cell death. None of the six candidates suppressed Zt6-induced cell death (Fig.4b,c), but one candidate, *PlLRR*, did suppress Zt12-induced cell death with potency similar to the *ZtLRR* control (Fig.4b,c). We aligned the predicted protein structure of each of these six PTI-suppressing sLRRs to ZtLRR, demonstrating a high level of conservation in the protein structure (highest: RccLRR, TM-score 0.88; lowest, MaLRR, TM-score 0.79) (Fig.S3). This indicates that structurally conserved sLRR proteins from diverse fungi have similar ability to suppress plant immune processes and that this phenomenon is not limited to *Z. tritici*.

**Figure 4.**
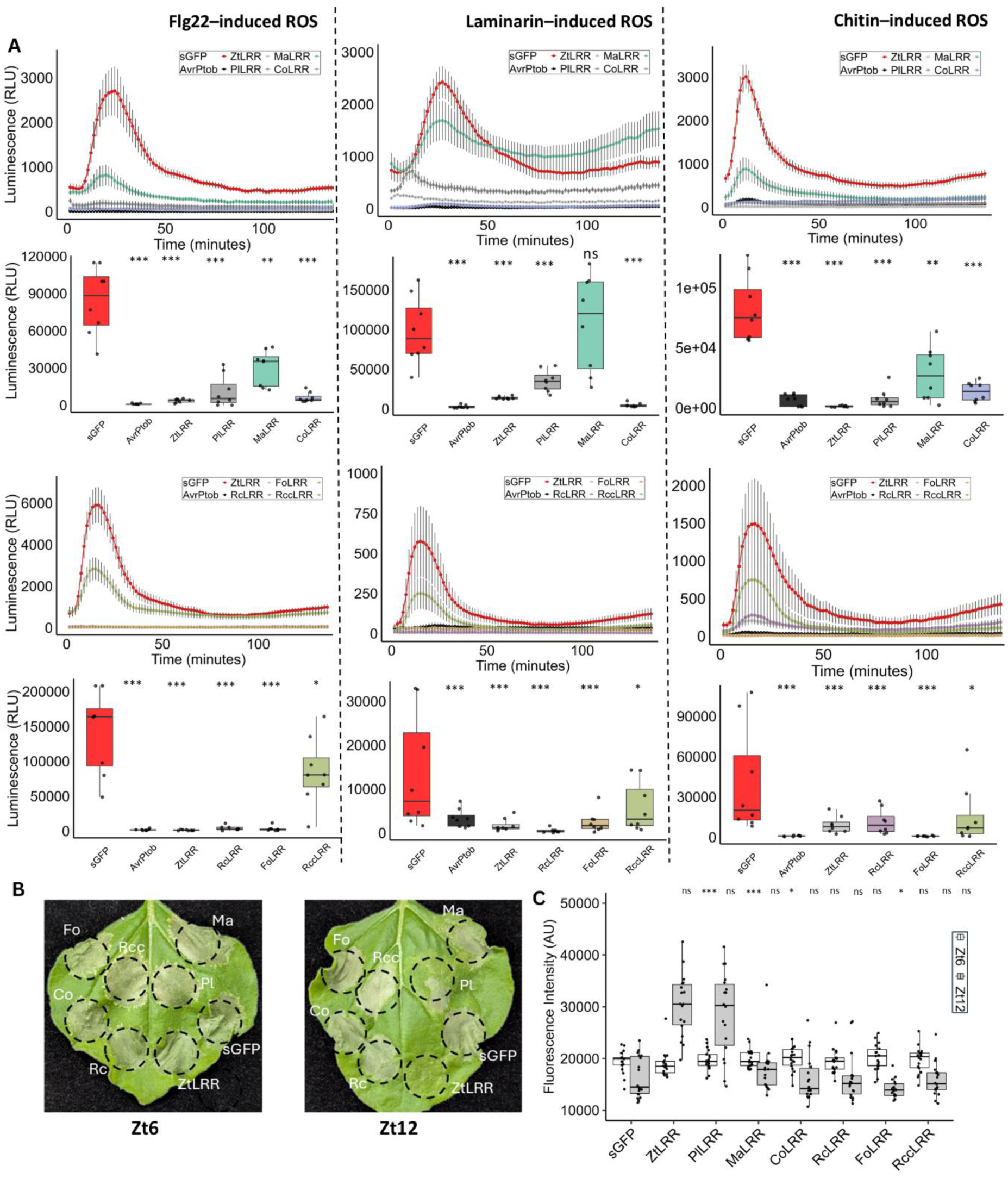
Structurally similar sLRR effectors from the Ascomycota and Mucoromycota suppress PTI and cell death in *N. benthamiana*. (A) Six candidate sLRR effectors from other fungi (PlLRR; *Plenodomus lindquistii*, MaLRR; *Mucor ambiguus*, CoLRR; *Colletotrichum orbiculare*, RcLRR; *Rhynchosporium commune*, FoLRR; *Fusarium oxysporum*, RccLRR; *Ramularia collo-cygni*) suppressed PAMP-induced ROS bursts in *N. benthamiana* induced by flg22 (upper), laminarin (middle) and chitin (lower panel). sGFP (red) and AvrPtoB/ZtLRR (dark grey/light grey) were used as controls. Asterisks indicate statistically significant differences at *p<0.05, **p<0.01, and ***p<0.001 from the sGFP control as performed by Tukey’s HSD test. (B) Of the six sLRRs from (A), only PlLRR was able to suppress Zt12-induced cell death, comparable to the ZtLRR control. Leaves were photographed at 5 dpi. (B) Quantification of Zt6- and Zt12-induced cell death suppression by the fungal sLRRs from (C). Boxplots show the red fluorescence intensity (arbitrary units) from infiltrated leaf patches. Statistical significance from the sGFP control was determined using Tukey’s HSD test; asterisks denote *P* < 0.05.

## Discussion

Using molecular mimics to hijack endogenous signalling pathways is an effective method of host manipulation by invading pathogens. Here we demonstrate that multiple fungal pathogens employ this technique with the use of sLRR proteins that are structural homologues of plant immune-related LRR-RLKs. Further, we show how these sLRRs can suppress both PTI and ETI to enhance virulence (Fig.5).

**Figure 5.**
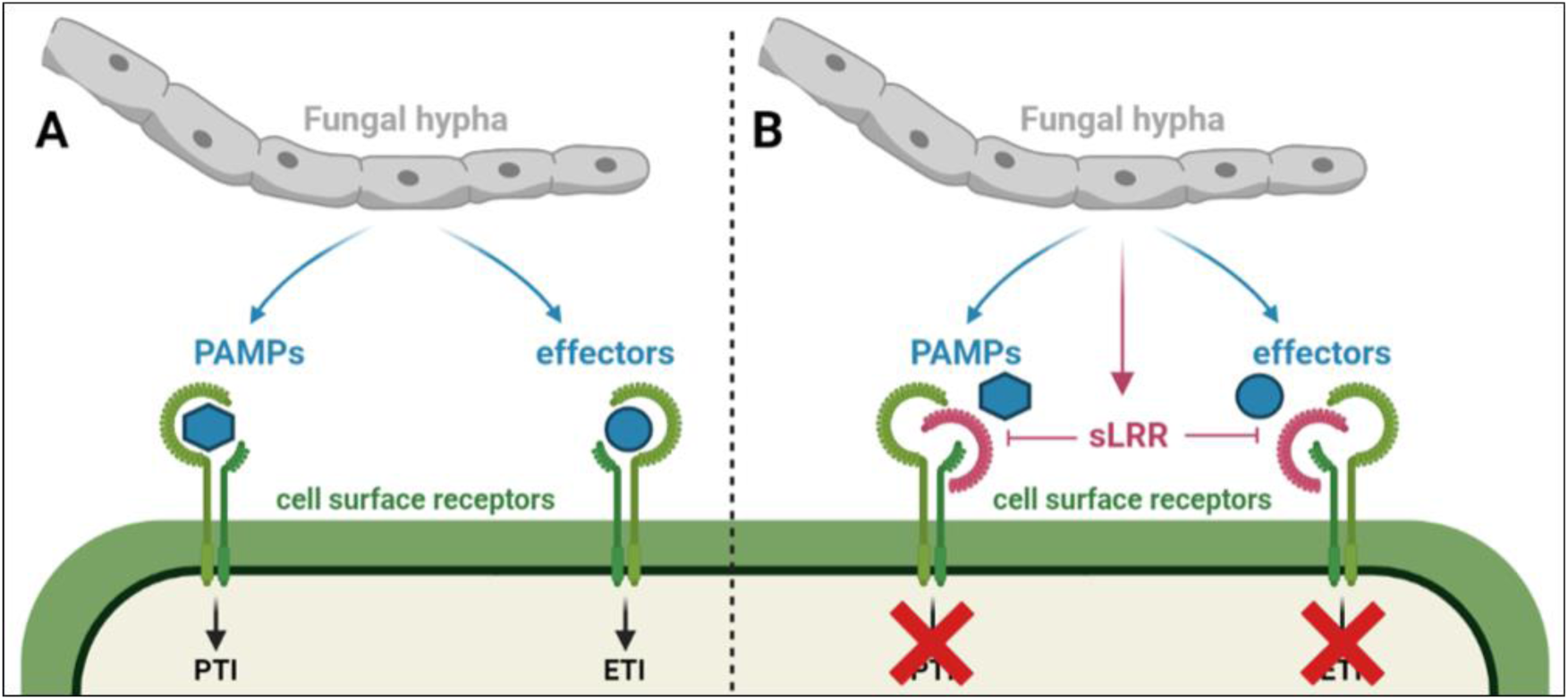
A model for sLRR-mediated immune suppression by receptor mimicry. (A) During invasive growth, hyphae shed PAMPs and secrete effectors which can be recognised by LRR-RLK immune complexes to trigger PTI/ETI. (B) sLRR effectors can interact with immune complex ectodomains to suppress PAMP/effector recognition and triggering of PTI/ETI.

Our focus organism is *Z tritici*, an interesting model for studying host-immune evasion due to its extended asymptomatic growth phase. The reference isolate for this fungus (IPO323) takes approximately two weeks after spore germination to induce necrotic lesions that are the hallmark of the disease septoria tritici blotch (STB). During this period, *Z. tritici* needs to evade immune surveillance to survive. However, once a compatible, virulent isolate of *Z. tritici* has initiated a successful infection, it establishes “systemic induced susceptibility” (SIS) (Seybold *et al*., 2020; Bernasconi *et al*., 2023). SIS suppresses the wheat immune system to a point where it enables independent avirulent isolates, to become virulent and commence infection (Bernasconi *et al*., 2023). Accordingly, SIS is thought to allow *Z. tritici* populations to maintain a diverse array of effectors and effector haplotypes (even in a single field population (Herreno *et al.,* 2025)) that may otherwise be lost due to selective pressure (Barrett *et al*., 2021; Suffert *et al*., 2024; Seybold *et al*., 2020; Bernasconi *et al*., 2023). The identification of ZtLRR as an immune suppressor, adds to a growing body of research describing immune-suppressive effectors expressed during wheat invasion (Thynne *et al*., 2024; Gomez-Gutierrrez *et al*., 2025). Indeed, this pathogen clearly has an extensive arsenal of effectors with immune-suppressive capability, that collectively, are essential to evade host defence for such a long period. Nonetheless, the mechanism(s) of SIS remain elusive, as our results demonstrate that ZtLRR is not responsible for inducing SIS. Although ZtLRR can suppress PTI triggered by three PAMPs (flg22, laminarin, and chitin) each of which signal through distinct receptors, constitutive expression of ZtLRR was not sufficient to overcome the gene-for-gene resistance mediated by the wall-associated kinase (WAK) resistance gene *Stb6* (Saintenac *et al*., 2018). The cognate effector partner of *Stb6*, *AvrStb6* (Zhong *et al*., 2017), is expressed in fungal hyphal tips during initial stomatal penetration (Alassimone *et al*., 2024). Accordingly, *Stb6*-mediated resistance occurs early during initial colonisation, prior to maximal expression of many of the known immune-suppressive effectors. We have previously speculated that if these suppressive effectors were induced earlier, they could assist in resistance breakdown; but this is not the case for ZtLRR and the AvrStb6-Stb6 interaction. However, constitutive expression of ZtLRR in wheat did enhance the virulence of an already compatible isolate, highlighting that this effector plays an important functional role in aiding infection development.

Signalling by cell-surface immune complexes is an important component of both PTI and ETI. BAK1 is required for signalling by numerous receptor complexes (Liebrand *et al*., 2014), therefore, it is not surprising it is frequently targeted by pathogen effectors. The bacterial effectors AvrPto, AvrPtoB and HopF2 directly interact with BAK1 to disrupt immune signalling (Shan *et al*., 2008; Zhou *et al*., 2014). The conserved fungal effector NIS1 can also interact with BAK1 and the associated receptor-like cytoplasmic kinase (RLCK) BIK1 to disrupt immunity (Irieda *et al*., 2019). These other effectors all function through interaction with the intracellular kinase domain of BAK1. To our knowledge, there is no other example of BAK1 targeting via the extracellular LRR domain. Therefore, ZtLRR and the other described fungal orthologs represent a novel mechanism for how pathogens can interfere with BAK1-mediated signalling from the apoplast. *Z. tritici* doesn’t form haustoria and it is unknown whether this fungus is able to deliver effectors intracellularly, where they might be able to access the BAK1 kinase domain or other proteins involved in immune signalling. The mimicry mechanism we propose here (Fig.5) may therefore be an adaptation to only having access to the ectodomains of BAK1 and associated receptors found in the apoplastic space.

A feature of ZtLRR and the other fungal sLRRs is the high level of predicted structural conservation, in contrast to the low level of primary sequence similarity and relatively disparate species distribution. However, despite this, we could still observe clear separation based on BLAST analyses, as to whether these fungal sLRRs were plant-like (i.e., more similar to clades of plant LRRs) or bacterial- and plant-like (i.e., more similar to bacterial and plant LRRs). Including ZtLRR, 7/11 plant-like sLRRs suppressed plant-immune responses. Despite structural conservation, only 1/5 of the bacterial sLRRs suppressed plant immune response. Our future work will investigate how subtle structural differences contribute to these variable phenotypes. For example, mutation of specific leucine residues may alter the overall fold conformation and lead to modified immune suppressing phenotypes. Another possibility is that the natural function of many sLRRs lies elsewhere, not related to suppression of ROS production or induction of cell death. This could include processes that are highly host-specific and not testable in the *N. benthamiana* model. Of the seven immune-suppressing sLRRs described in this study (including ZtLRR), six are from Ascomycete species (two Sodariomycetes, three Dothideomycetes, one Leotiomycete). All of these species are hemibiotrophic phytopathogens, therefore sLRRs may be an evolutionary innovation associated with this lifestyle. Each of these sLRRs belong to the “plant-like sLRR” clade. The remaining immune-suppressing sLRR is from *Mucor ambiguus*, a Mucoromycete. This sLRR belongs to the "bacterial-like" clade of sLRRs, and was able to suppress ROS production induced by two of the three tested PAMPs. Given that the Mucoromycota and the Dikarya are thought to have diverged >600 Mya (Zhao *et al.,* 2023), this raises the intriguing question of whether sLRR-mediated molecular mimicry has (i) evolved independently in this lineage, or (ii) is ancient, yet only conserved in a small number of species where it offers selective advantage. Our future work will aim to answer this question.

The highly conserved structures of the fungal sLRRs indicates a selective pressure in the harbouring species to maintain highly similar conformation to their plant LRR protein counterparts. Accordingly, these fungal sLRRs likely interact with similar proteins or molecules to their plant counterparts. Plant homologues include the ectodomains of RLKs and receptor-like proteins (RLPs), and leucine-rich extensin-like proteins (LRXs). Our hypothesis is that fungal sLRRs mimic LRR structures in receptor ectodomains to inhibit or otherwise interfere with the interaction of RLKs and their conserved network-hub, which often includes the co-receptor BAK1 (Fig.5). Our yeast two-hybrid data (Fig.2) suggest this to be a likely mode of action. This could explain how several fungal sLRRs are able to suppress flg22-induced immune responses, as this bacterial PAMP is perceived by a FLS2/BAK1 complex. However, this does not explain how these effectors suppress laminarin- and chitin-induced PTI, which are recognised in a BAK1-independent manner by CERK receptors (chitin) or through unknown mechanisms (laminarin). It is possible that ZtLRR and other sLRRs are relatively promiscuous proteins, able to interact with a range of cell-surface receptors, many of which remain to be identified. Future work could utilise screen-based approaches to identify all possible ZtLRR interactors in the apoplastic environment. Another possibility is that fungal sLRRs function similarly to the endogenous protein ligands of LRR-RLKs that are negative regulators of defence receptor signalling. Plant pathogens can use additional molecular mimic proteins such as CEP, CLE and RALF peptides. Plant LRX proteins interact with RALF peptides to coordinate their function, so perhaps fungal sLRRs play a similar role.

## Acknowledgements

HA was funded by a PhD studentship award from the Darwin Trust of Edinburgh with GK as supervisor. ET was funded by a Marie Skłodowska-Curie Actions fellowship. RB, MSH and EW were funded by BBSRC grant BB/R014876/1, A Community Resource for Wheat and Rice Transformation. IMLS and EEC were funded by the DFG Emmy Noether Programme (SA 4093/1-1 to IMLS) from the DFG (Deutsche Forschungsgemeinschaft/German Research Foundation) and acknowledge support from the Cluster of Excellence on Plant Sciences (CEPLAS) funded by the DFG under Germany’s Excellence Strategy–EXC 2048/1 (Project ID: 390686111) and the DFG Collaborative Research Centre (Project ID: 414786233; SFB 1403 to IMLS). ET would like to acknowledge Ksenia Krasileva (University of California, Berkeley) for helpful discussions early in this project.

## Author contributions

GK and ET initiated the project. GK and EHS financially supported the project. HA did the *N. benthamiana* and wheat ROS assays, and the wheat pathoassays. GK performed the *N. benthamiana* cell death experiments and generated the ZtLRRΔSP construct. TE identified the sLRR orthologs and performed the related cloning procedures. CB did qPCR using ZtLRR transgenic wheat and performed genotyping. LP performed protein structure predictions and structural phylogenetic analyses. EC and IMLS generated wheat protoplasts and performed the protoplast ROS assays. EW, RB, MSH generated the transgenic wheat lines. GK, ET and HA wrote the manuscript.

## Supplementary material

**Supplementary Figure 1. Plant-like sLRR homologues are present among various Dothideomycete, Sordariomycete, and Leotiomycete species.** Representation of a species tree, broadly adapted from the Mycocosm (https://mycocosm.jgi.doe.gov) species trees of fungal lineages. Species with plant-like LRR homologues are labelled in blue. Representative species from different fungal families and genera that lack LRR homologues are labelled in black. Species labelled in purple (*Setosphaeria turcica*) represents a species with a plant- and bacterial-like LRR.

**Supplementary Figure 2. Structure-based phylogeny of fungal and oomycete sLRRs, compared to related plant and bacterial LRRs. L**RR structures predicted with AlphaFold3. Fungal LRR (tan), Oomycete LRRs (light green), Plant LRRs (dark green), and bacterial LRRs (peach).

**Supplementary Figure 3. Protein model alignments of PTI-suppressors with ZtLRR demonstrate high levels of structural conservation among sLRRs.** The structures of each of the six PTI-suppressing sLRRs were predicted using AlphaFold3, with high levels of confidence (highest: PlLRR, pTM 0.94; lowest: MaLRR, pTM 0.74). Protein alignments with ZtLRR demonstrate high levels of conservation of predicted structure among these sLRRs (highest: RccLRR, TM-score 0.88; lowest, MaLRR, TM-score 0.79).

**Supplementary Table 1. Summary of 15 fungal/oomycete sLRR orthologs cloned for functional characterisation.**

**Supplementary File 1. BLAST hits (150) for each plant-like sLRR (BLASTp; NCBI-NR database).**

**Supplementary File 2. BLAST hits (150) for each bacterial and plant-like sLRR (BLASTp; NCBI-NR database).**

