## Supplementary Figures for "Molecular mimicry of plant cell-surface immune receptors by fungal secreted leucine-rich repeat proteins"

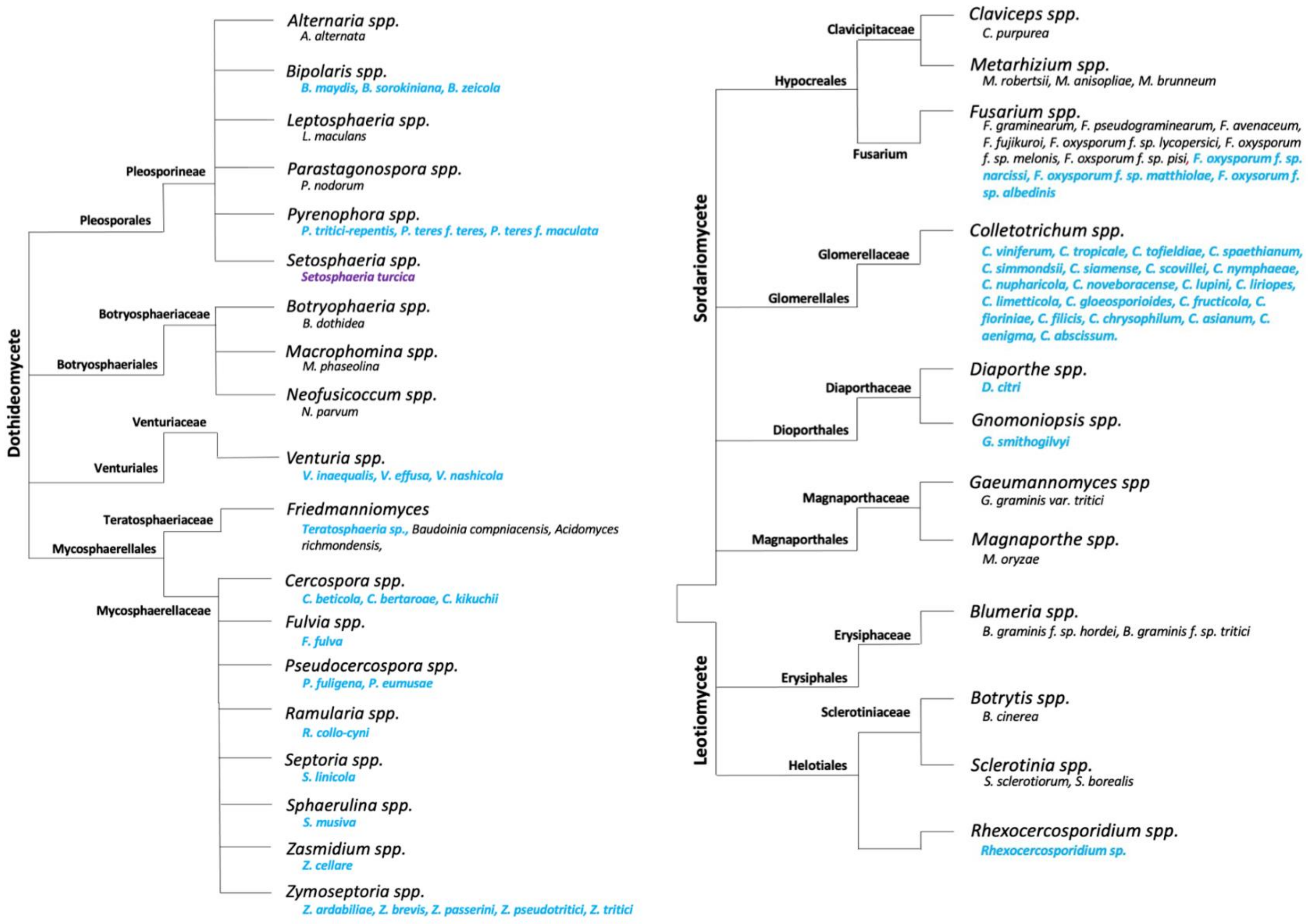


**Supplementary Figure 1. Plant-like sLRR homologues are present among various Dothideomycete, Sordariomycete, and Leotiomycete species.** Representation of a species tree, broadly adapted from the Mycocosm (https://mycocosm.jgi.doe.gov) species trees of fungal lineages. Species with plant-like LRR homologues are labelled in blue. Representative species from different fungal families and genera that lack LRR homologues are labelled in black. Species labelled in purple (*Setosphaeria turcica*) represents a species with a plant- and bacterial-like LRR.

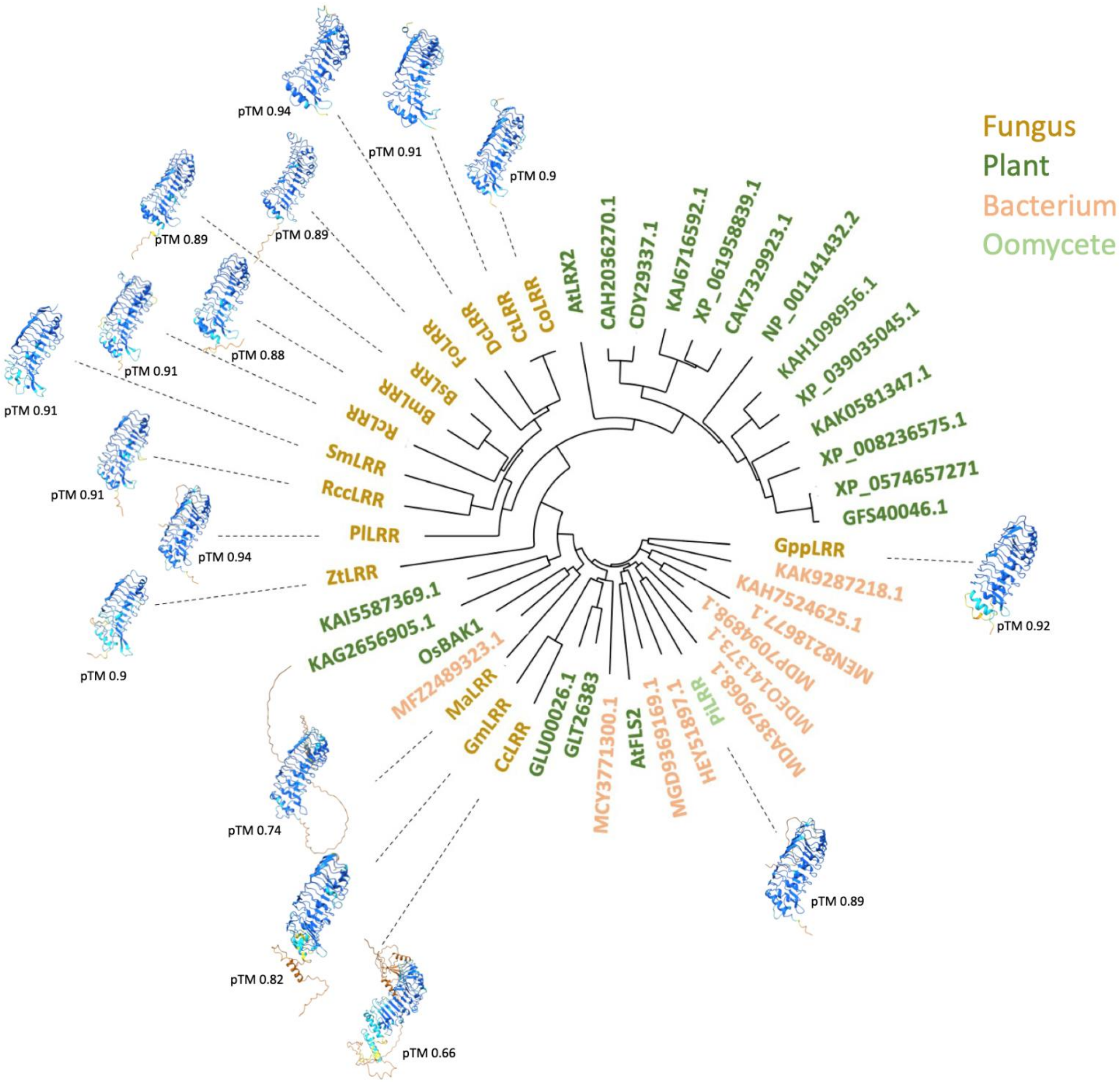


**Supplementary Figure 2. Structure-based phylogeny of fungal and oomycete sLRRs, compared to related plant and bacterial LRRs. L**RR structures predicted with AlphaFold3. Fungal LRR (tan), Oomycete LRRs (light green), Plant LRRs (dark green), and bacterial LRRs (peach).

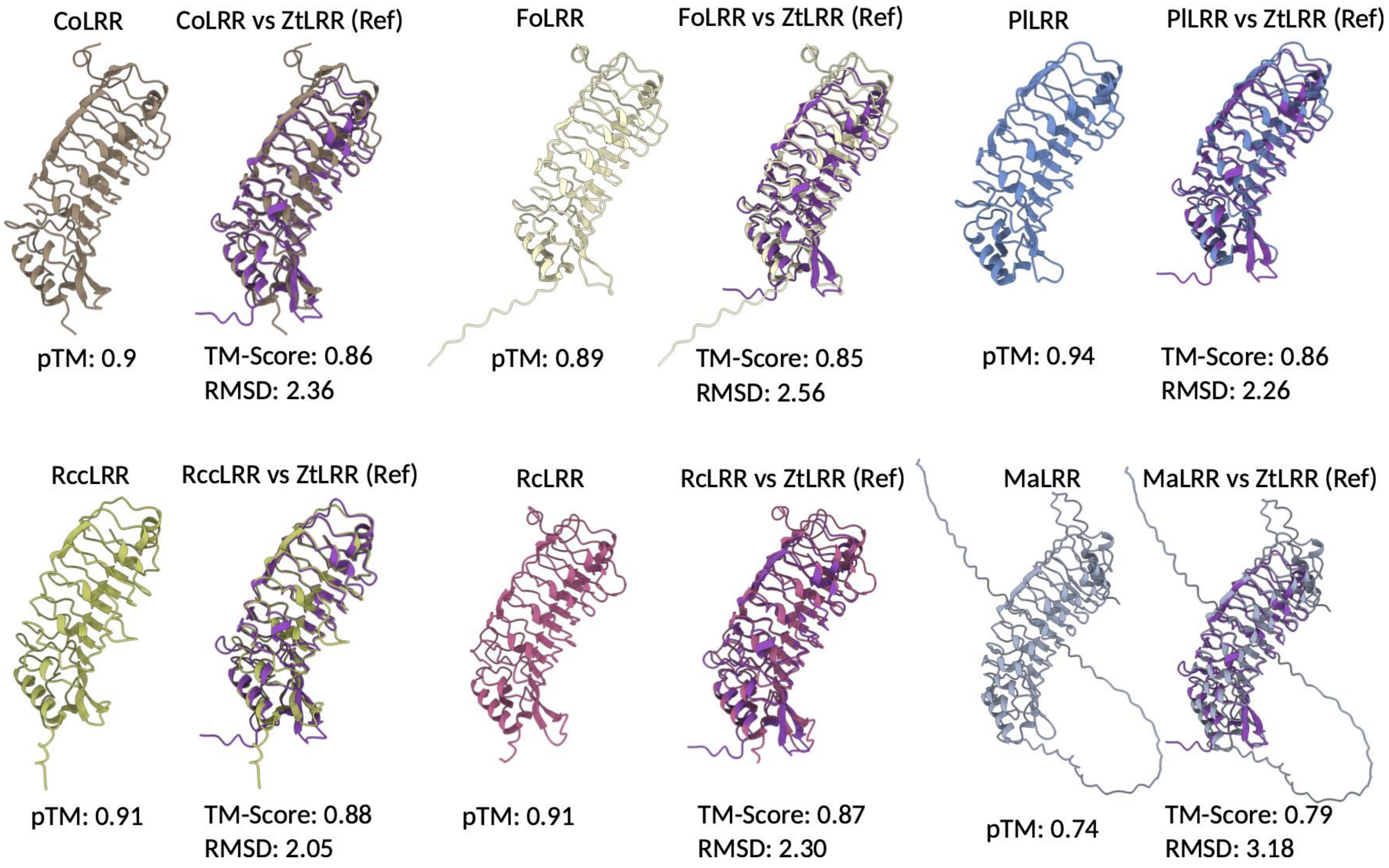


**Supplementary Figure 3. Protein model alignments of PTI-suppressors with ZtLRR demonstrate high levels of structural conservation among sLRRs.** The structures of each of the six PTI-suppressing sLRRs were predicted using AlphaFold3, with high levels of confidence (highest: PlLRR, pTM 0.94; lowest: MaLRR, pTM 0.74). Protein alignments with ZtLRR demonstrate high levels of conservation of predicted structure among these sLRRs (highest: RccLRR, TM-score 0.88; lowest, MaLRR, TM-score 0.79).
